# Real Time Normalization of Fast Photochemical Oxidation of Proteins Experiments by Inline Adenine Radical Dosimetry

**DOI:** 10.1101/352385

**Authors:** Joshua S. Sharp, Sandeep K. Misra, Jeffrey J. Persoff, Robert W. Egan, Scot R. Weinberger

## Abstract

Hydroxyl radical protein footprinting (HRPF) is a powerful method for measuring protein topography, allowing researchers to monitor events that alter the solvent accessible surface of a protein (e.g. ligand binding, aggregation, conformational changes, etc.) by measuring changes in the apparent rate of reaction of portions of the protein to hydroxyl radicals diffusing in solution. Fast Photochemical Oxidation of Proteins (FPOP) offers an ultra-fast benchtop method for performing HRPF, photolyzing hydrogen peroxide using a UV laser to generate high concentrations of hydroxyl radicals that are consumed on roughly a microsecond timescale. The broad reactivity of hydroxyl radicals means that almost anything added to the solution (e.g. ligands, buffers, excipients, etc.) will scavenge hydroxyl radicals, altering their half-life and changing the effective radical concentration experienced by the protein. Similarly, minute changes in peroxide concentration, laser fluence, and buffer composition can alter the effective radical concentration, making reproduction of data challenging. Here, we present a simple method for radical dosimetry that can be carried out as part of the FPOP workflow, allowing for measurement of effective radical concentration in real time. Additionally, by modulating the amount of radical generated, we demonstrate that FPOP HRPF experiments carried out in buffers with widely differing levels of hydroxyl radical scavenging capacity can be normalized on the fly, yielding statistically indistinguishable results for the same conformer. This method represents a major step in transforming FPOP into a robust and reproducible technology capable of probing protein structure in a wide variety of contexts.

## Introduction

Hydroxyl radical protein footprinting (HRPF) is an emerging technology for structural biology that offers protein topographical information down to single amino acid side chain resolution. While HRPF does not offer the structural resolution of X-ray crystallography or multidimensional NMR spectroscopy, it has the noted advantage of using liquid chromatography coupled to tandem mass spectrometry (LC-MS/MS) as its measurement technique, offering greater flexibility in sample types and solution conditions that can be probed.^*1*^ The underlying principle of HRPF is that the rate of oxidation of a given amino acid is based on two factors: the inherent reactivity of the amino acid (which depends on the structure of the side chain^*2*^ and the sequence context of the amino acid^*3, 4*^) and the accessibility of the side chain to the hydroxyl radical diffusing in solution^*3, 5–7*^. Samples are exposed to a concentration of hydroxyl radicals generated by any of a variety of means^*8–17*^. Hydroxyl radicals rapidly react with amino acid side chains, ultimately forming stable modification products that are dominated by the net addition of one or more oxygen atoms^*2, 18–20*^. These modifications are stable, freezing information about protein topography in a chemical “snapshot” taken at the time of radical exposure; therefore, samples can be processed after oxidation and quenching by any of a variety of means including denaturation and reduction of disulfide bonds, proteolysis, and deglycosylation. By measuring changes in the apparent rate of oxidation of a given amino acid or stretch of amino acids by LC-MS/MS as a protein goes from one structural state to another (e.g. ligand-bound vs. ligand free, monomer vs. aggregate, etc.), changes in the topography of the protein or protein complex can be measured.

Fast Photochemical Oxidation of Proteins (FPOP) is a method for generating hydroxyl radicals for HRPF. In FPOP, the protein sample of interest is mixed with hydrogen peroxide, along with a radical quencher to limit radical half-life (e.g. glutamine). This mixture is pushed through a fused silica capillary into the path of a focused UV laser (usually a KrF excimer laser or a frequency quadrupled Nd:YAG). The pulse rate of the laser and the illuminated length of the capillary are chosen, along with the flow rate of the solution through the capillary, to ensure that each volume of sample is illuminated by a single laser pulse, with an exclusion volume maintained to correct for laminar flow and sample diffusion^*14*^. The sample is then deposited into a quenching solution containing catalase and methionine amide to quickly eliminate excess hydrogen peroxide, secondary oxidants like superoxide, and protein hydroperoxides^*14, 21*^. FPOP has several advantages that have made it a very popular choice: it is a benchtop method that can be performed in a variety of labs; it generates a very short burst of hydroxyl radicals that are consumed very quickly (on a microsecond time scale), allowing the initial radical attack to be completed faster than the protein can unfold due to the labeling event^*22*^; it is capable of very high levels of label density^*23*^; it is compatible with pulse-probe experimental designs, allowing it to make time-sensitive measurements^*24–26*^; and it is flexible regarding sample, allowing for measurements of integral membrane proteins^*27–33*^ and even intact cells^*34, 35*^.

However, like all covalent labeling technologies that use a broadly reactive label, HRPF by FPOP is very sensitive to changes in the solution. Adding or changing the concentration of almost any organic reagent (e.g. buffer, ligand, detergent, lipids, excipients, allosteric modulator, etc.) offers a new competing pathway for consumption of hydroxyl radicals, changing the half-life of the radical and altering the effective concentration of hydroxyl radical experienced by the protein.

Additionally, changes in laser fluence, hydrogen peroxide concentration, or even hydrogen peroxide age can significantly alter the effective concentration of hydroxyl radical, making reproducibility and robustness of HRPF by FPOP a major problem. Several options for measuring effective radical concentrations in FPOP have been published, including peptide-based radical dosimetry^*23, 36*^ and isotope dilution GC-MS^*37*^. Perhaps the simplest was to replace a portion of the glutamine radical scavenger with adenine, which has an almost identical rate of reaction as glutamine^*38*^. Adenine does not change its UV absorbance in the presence of hydrogen peroxide or after UV illumination from the laser; however, upon reaction with hydroxyl radicals, the UV absorbance of adenine around 260 nm decreases substantially. Xie and Sharp previously showed that monitoring the UV absorbance of the adenine in an FPOP reaction after quenching measures the effective hydroxyl radical concentration, and allows the researcher to ensure that two samples experienced equivalent FPOP conditions for comparison^*39*^. Regardless of the method, post-quenching radical dosimetry is quickly becoming a standard part of best practices for any FPOP experimental protocol^*36*^.

While post-quenching dosimetry offers a method for monitoring effective hydroxyl radical concentration, normalization of samples with different scavenging capacities is cumbersome. The researcher must perform FPOP and quenching of the sample under each condition, then measure the dosimeter response. If the dosimeter response was not equivalent, the researcher must estimate the amount of additional radical generation required to generate an equivalent response and repeat the experiment, modifying conditions until an equivalent response is generated. This process is tedious and wasteful of sample. Here, we present a method for inline hydroxyl radical dosimetry that measures effective radical dose in real time, immediately after illumination with the UV laser and prior to quenching. This inline radical dosimetry is achieved through the use of a high efficiency UV absorbance detector that continuously measures UV absorbance at 265 nm, monitoring the absorbance of adenine immediately after oxidation^*39*^. Using this inline radical dosimetry system, we demonstrate that differential radical scavenging capacities for samples can be compensated in real time, allowing for efficient experimental normalization and significantly improving the convenience, reproducibility and robustness of FPOP HRPF.

## Experimental

### Materials

[Glu]_1_-Fibrinopeptide B (GluB), myoglobin, catalase, formic acid, glutamine, sodium phosphate, 2-(N-morpholino)-ethanesulfonic acid (MES), and sequencing grade modified trypsin were obtained from Sigma-Aldrich (St. Louis, MO, USA). Adenine and LCMS-grade acetonitrile and water were purchased from Fisher Scientific (Fair Lawn, NJ, USA). Hydrogen peroxide (30%) was purchased from J. T. Baker (Phillipsburg, NJ, USA). Fused silica capillary was purchased from Molex, LLC (Lisle, IL, USA).

### FPOP and Inline Dosimetry

The peptide and protein normalization samples used a final concentration of 2 μM of GluB and 5 μM of myoglobin mixed together. Samples were mixed to a final concentration of 100 mM hydrogen peroxide (unless otherwise noted), 16 mM glutamine, and 2 mM adenine immediately prior to illumination. Samples were loaded into a 100 μL gastight syringe (Hamilton, Reno, NV, USA) and mounted on a Legato 101 syringe pump (KD Scientific, Holliston, MA, USA) with a calculated exclusion volume of 15%, under FPOP module component software control (GenNext Technologies, Montara, CA). The sample was pushed through a 100 μm ID, 150 μm OD fused silica capillary through the beam path of a COMPex Pro 102 KrF excimer laser (Coherent Inc., Santa Clara, CA, USA) focused with a convex lens (Edmund Optics, Barrington, NJ, USA). The polyimide coating of the fused silica capillary was burned off by pre-illumination with the UV laser beam until the capillary was free of the coating at the site of laser illumination by visual inspection. Immediately after the illumination window, the sample was pushed through the inline radical dosimeter (GenNext Technologies) and measured as described below. Immediately after dosimetry, sample was deposited into an automated product collector module (GenNext Technologies) that deposited sample either into waste or into a quench solution of 0.5 μg/μL methionine amide and 0.5 μg/μL catalase to eliminate secondary oxidants.

### LC-MS Analysis

50 mM Tris, pH 8.0 and 1 mM CaCl_2_ was added to the myoglobin and GluB samples after FPOP. The sample was incubated at 80o C for 20 minutes to denature the protein. After the sample had been cooled to room temperature, a 1:20 trypsin:protein weight ratio was added to the samples for overnight digestion at 37 ° C with sample rotation. Digestion was terminated by heating the samples to 95° C for 10 min.

The protein and peptide samples were analyzed on an Orbitrap Fusion instrument controlled with Xcalibur version 2.0.7 (Thermo Fisher, San Jose, CA). Samples were loaded onto an Acclaim PepMap 100 C18 nanocolumn (0.75 mm × 150 mm, 2 μm, Thermo Fisher Scientific). Separation of peptides on the chromatographic system was performed using mobile phase A (0.1% formic acid in water) and mobile phase B (0.1% formic acid in acetonitrile) at a rate of 300 μL/min. The peptides were eluted with a gradient consisting of 2 to 35% solvent B over 22 min, ramped to 95 % B over 5 min, held for 3 min, and then returned to 2% B over 3 min and held for 9 min. Peptides were eluted directly into the nanospray source of an Orbitrap Fusion instrument using a conductive nanospray emitter (Thermo Scientific). All data were acquired in positive ion mode. The spray voltage was set to 2300 volts, and the temperature of the heated capillary was set to 300° C. In CID mode, full MS scans were acquired from m/z 350 to 2000 followed by eight subsequent MS/MS scans on the top eight most abundant peptide ions.

### HRPF Data Analysis

Unoxidized and oxidized peptide peaks were identified by LC-MS and LC-MS/MS using ByOnic version v2.10.5 (Protein Metrics, San Carlos, CA, USA). Samples were quantified by integration of the selected ion chromatogram peaks of peptides plus one or more oxygen atoms (mass error = 10 ppm), with all resolved oxidation isomers summed. Oxidation events per peptide were calculated using equation 1: 
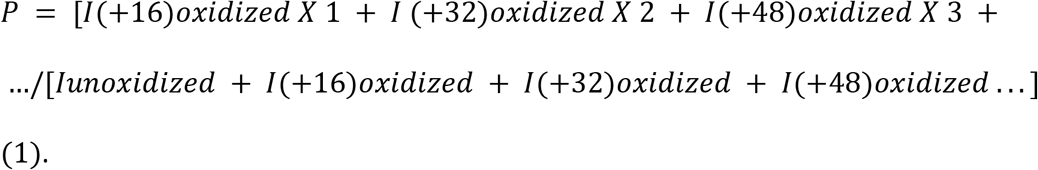
where *P* denotes the oxidation events at the peptide level and *I* values are the peak intensities of oxidized and unoxidized peptides. Unilluminated control samples were used to measure background oxidation in the absence of laser UV light. These levels were very low, and when detected were subtracted from the measured FPOP oxidation.

### Inline Dosimeter Design

The radical dosimeter was placed inline after the illumination zone of the fused silica capillary, prior to sample collection and quenching. Immediately after the illumination region, the illumination capillary (100 μm ID, 150 μm OD) was terminated in a low dead volume union, where it was joined with a 100 μm ID, 362 μm OD fused silica capillary within which the dosimetry would take place. The illumination capillary was separated from the dosimetry capillary to allow for replacement of the illumination capillary in the event of laser-induced breakage without requiring replacement of the dosimetry capillary. An ultraviolet light detection window was formed in the dosimetry capillary by burning off the polyimide coating for approximately a 1 cm axial length. The dosimetry capillary was then installed into the inline radical dosimeter, within a capillary holder that aligns the detection window with incident 265 nm light produced by a UV light emitting diode (bandwidth +/− 6 nm). Current from the photodetection of transmitted light within the radical dosimeter was converted to voltage by a low noise trans-impedance amplifier. Output voltage was subsequently converted to a 16-bit digital signal and absorbance measurements ultimately calculated by the system’s custom software program.

The inline radical dosimeter optical bench is depicted in **Figure 1**. Light was focused into the capillary lumen by a quartz ball lens. Transmitted light was collimated by a second ball lens and directed to strike a UV-sensitive silicon photodiode. The probed axial length of the capillary was 0.1 mm, making the probed volume of a 100 μm ID capillary 0.8 nL. During operation, the decrease in adenine absorbance at 265 nm (ΔAbs_265nm_) was determined by comparing the absorbance of an unirradiated sample with that of a laser-irradiated sample. Immediately after the inline dosimetry unit, the sample was automatically deposited into a quenching solution held in an FPOP HRPF product collector.

**Figure 1.**
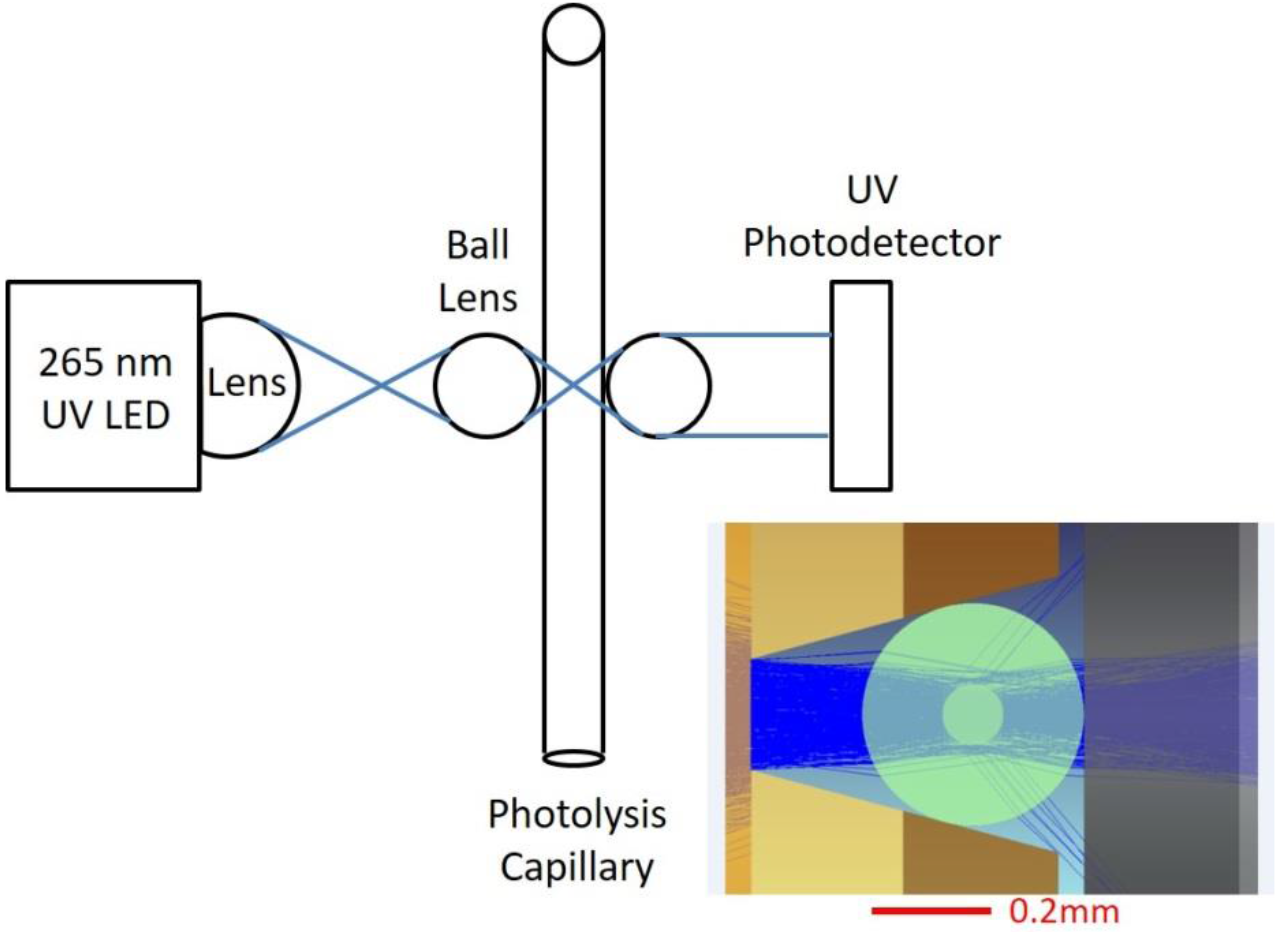
Optical layout of the inline dosimeter. A V-groove positions a 362 μm OD capillary precisely at the optimal focal plane for the UV LED. Inset: Modeled light path for the UV LED. The vast majority of the light passes through the lumen of the capillary into the detector.

### Dosimetry System Software, HRPF Product Collector, and FPOP Component Control Module

FPOP System Control and Dosimetry Data Analysis Software, working in concert with an FPOP Component Control Module (GenNext Technologies) and an HRPF Product Collector, provided for control and coordination all FPOP system components, as well as for the collection, display and processing of the inline dosimeter UV signal.

The Control Module interfaced with the FPOP system via a USB, and with the Legato 101 syringe pump (KD Scientific, Holliston, MA, USA) using a TTL output. The pump flow rate, laser repetition rate and the start/stop of the laser were under control of the respective instrument software/firmware.

The dosimeter software displayed real-time readings of UV absorbance based on the voltage from the inline dosimeter, with both auto-zeroing and manual zeroing supported. The inline dosimeter UV voltage signal was read using a 16-bit differential analog input at 1 KHz with 100 points averaged, then converted to give a 10 Hz effective absorbance data rate. A user selectable (0-5 second) simple moving average filter was applied for absorbance data smoothing. The software also calculated the post-dosimeter residence time based on user inputs of capillary ID, post-dosimeter capillary length, and syringe pump flow rate. Upon user input that the desired adenine absorbance had been reached, the dosimeter software would calculate the sample residence time in the post-dosimeter capillary length based on the post-dosimeter capillary length, ID, and flow rate. After this time was reached, the software would direct the Product Collector to move from a waste vial to a product collection vial to ensure that only sample exposed to the desired effective concentration of hydroxyl radical was collected.

## Results and Discussion

### Inline Dosimetry Response

We tested the linearity of response of the inline radical dosimeter to increasing levels of hydroxyl radical concentration. We generated hydroxyl radicals in the presence of 2 mM adenine, 16 mM glutamine, and water at concentrations of hydrogen peroxide ranging from 20 mM to 100 mM, at a laser fluence of 9.08 mJ/mm^2^. As shown in **Figure 2**, ΔAbs_265nm_ varied linearly, increasing as the hydrogen peroxide concentration (and, therefore, the amount of hydroxyl radical generated) increased. The coefficient of variation was low and decreased as the radical concentration increased, ranging from 0.12 at 20 mM peroxide to 0.017 at 100 mM peroxide.

**Figure 2.**
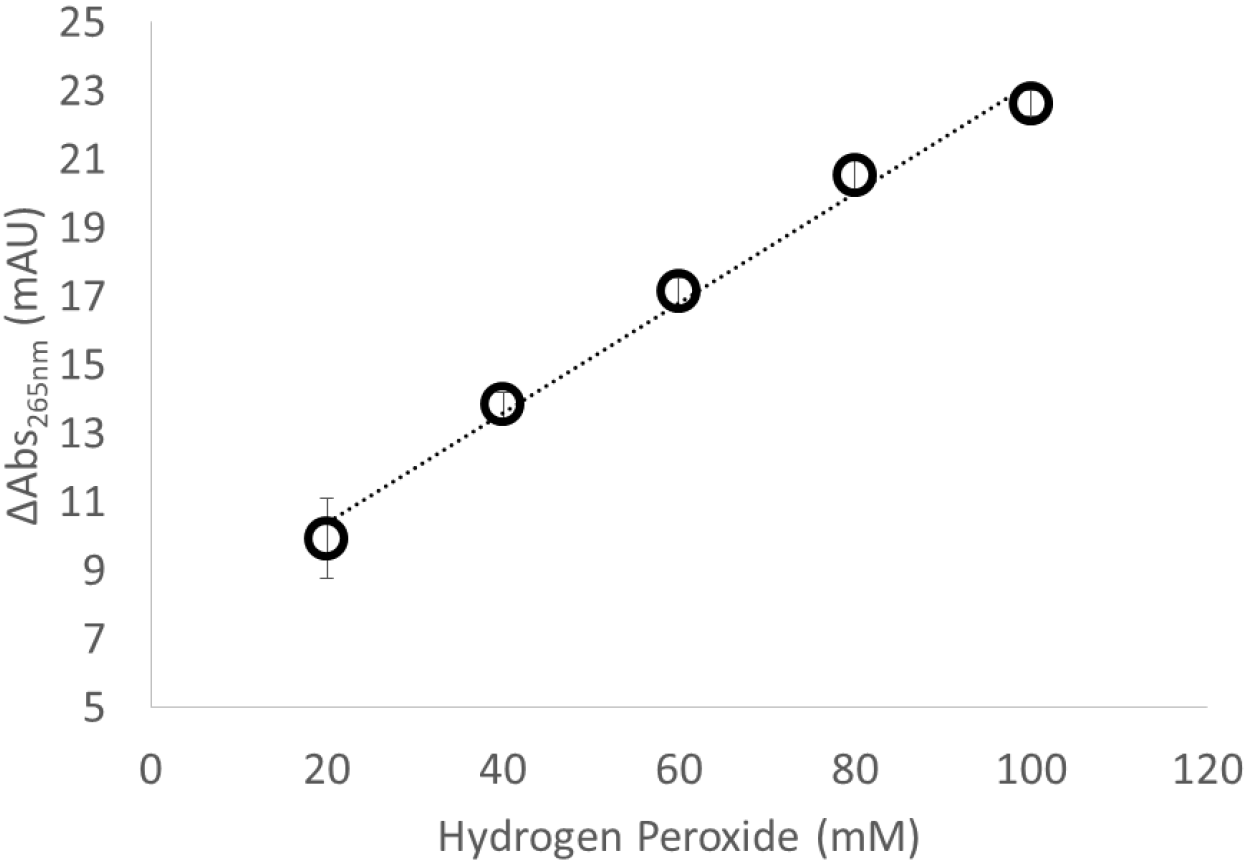
Change in adenine absorbance as a function of hydrogen peroxide concentration. As the amount of hydroxyl radical generated increases, the dosimeter response increases linearly.

We also tested the linearity of response to changes in the radical scavenging capacity of the solution. We generated hydroxyl radicals in the presence of 2 mM adenine, 16 mM glutamine, 100 mM hydrogen peroxide, and water at concentrations of MES buffer (an organic buffer that is a very efficient hydroxyl radical scavenger) ranging from 1 mM to 10 mM, at a laser fluence of 9.50 mJ/mm2. As shown in **Figure 3**, the inline radical dosimeter is able to detect the decrease in the effective hydroxyl radical concentration due to the increased scavenging capacity of the MES buffer, with ΔAbs_265nm_ decreasing linearly with MES buffer concentration. Precision was again good, with the coefficient of variation ranging from 0.14 to 0.05, although in this case the change in precision did not track directly with dosimeter response. Interestingly, we detected an anomalous response from Tris buffers; when oxidized, absorbance at 265 nm increases (**Figure S1**, Supporting Information), confounding standard adenine dosimetry.

**Figure 3.**
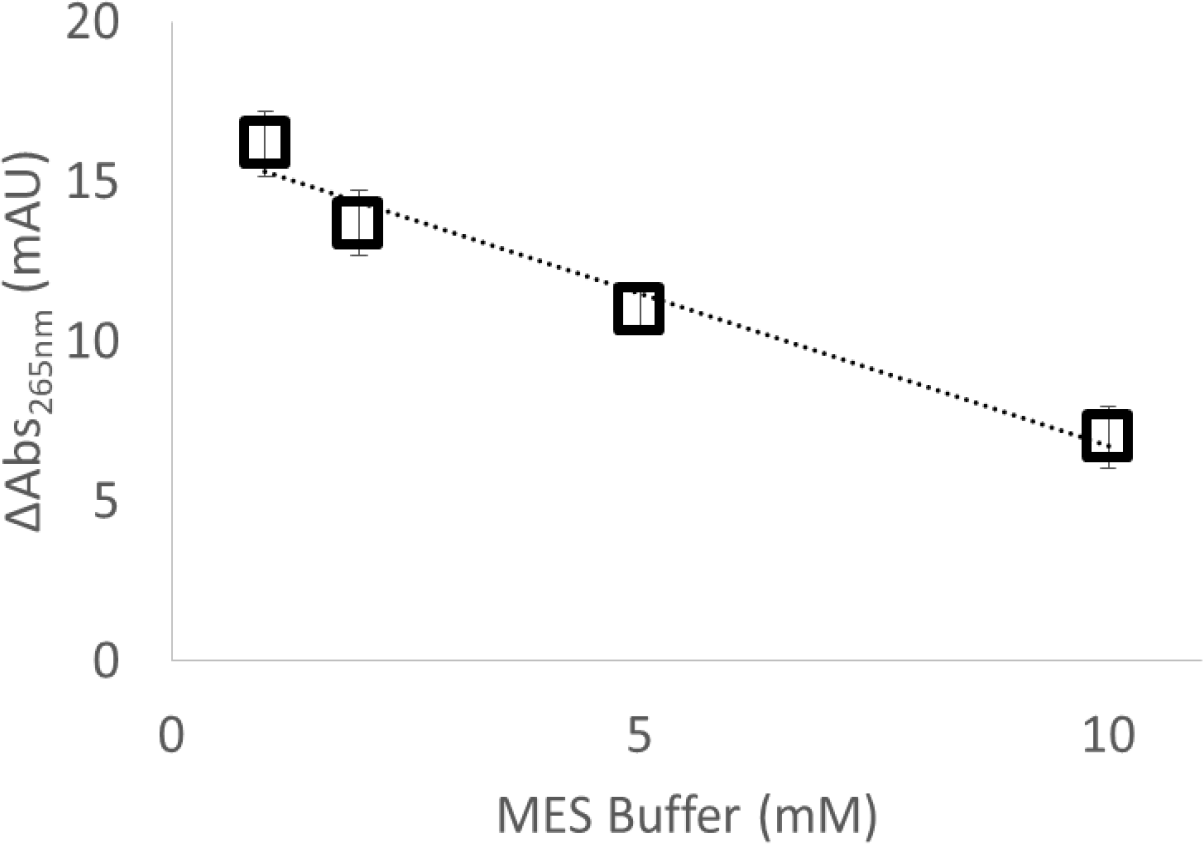
Change in adenine absorbance as a function of radical scavenger concentration. As the amount of MES radical scavenger increases, the effective radical dose decreases and the dosimeter response decreases linearly.

### Inline Dosimetry-Based Normalization of FPOP

In order to determine if inline adenine dosimetry is capable of correcting for differential sample scavenging capacities in real time, we analyzed 2 μM GluB and 5 μM myoglobin in 5 mM sodium phosphate buffer (pH, 7.4 X) by FPOP in triplicate at laser fluence of 13.81 mJ/mm2 with a 15% exclusion volume. These analyses yielded a ΔAbs_265nm_ of 13.93 ± 2.37 mAU. We then analyzed the same concentration of GluB and myoglobin in 5 mM MES buffer (pH, 7.4) by FPOP, adjusting the laser energy per pulse until we approached an identical ∆Abs_265nm_ (13.27 ± 0.29 mAU) at laser fluence of 15.1 mJ/mm2. Flow rate was adjusted to maintain a 15% exclusion volume, as the higher laser energy altered the beam area. After equivalent ΔAbs_265nm_ was achieved at the dosimeter, the sample was collected and quenched for analysis. After quenching and digestion, the oxidation of GluB was measured by LC-MS and shown in **Figure 4**. The two samples are not statistically different (*p* = 0.39). These results indicate that online dosimetry allows for real time adjustment of FPOP conditions to overcome differential scavenging and experimental conditions.

**Figure 4.**
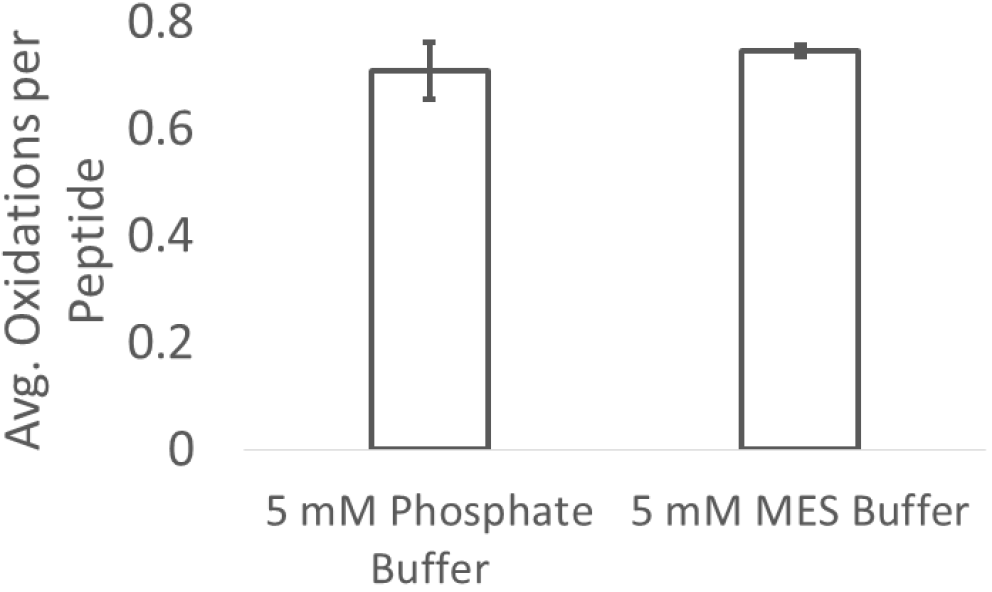
Real time normalization of GluB oxidation by online adenine dosimetry. GluB oxidized in phosphate buffer at 13.81 mJ/mm^2^ yielded a ΔAbs_265nm_ of 13.93 ± 2.37 mAU. For GluB in MES buffer, laser energy was adjusted until an equivalent ΔAbs_265nm_ was achieved (13.27 ± 0.29 mAU). Both samples gave equivalent levels of peptide oxidation as measured by LC-MS.

While **Figure 4** shows that we can normalize FPOP in real time by online adenine dosimetry to adjust for differential sample scavenging capacity using a standard reporter peptide assay, the most common application would be to normalize protein footprints from different experimental conditions and/or laboratories. We also tested the ability of online adenine dosimetry to normalize the hydroxyl radical footprint of myoglobin. As observed in **Figure 5**, myoglobin gave footprinting results that are not statistically different under two very different radical scavenging conditions. These data clearly demonstrate that FPOP experiments can be normalized in real time under very different experimental conditions by employing inline adenine dosimetry, while requiring minimal use of time and sample.

**Figure 5.**
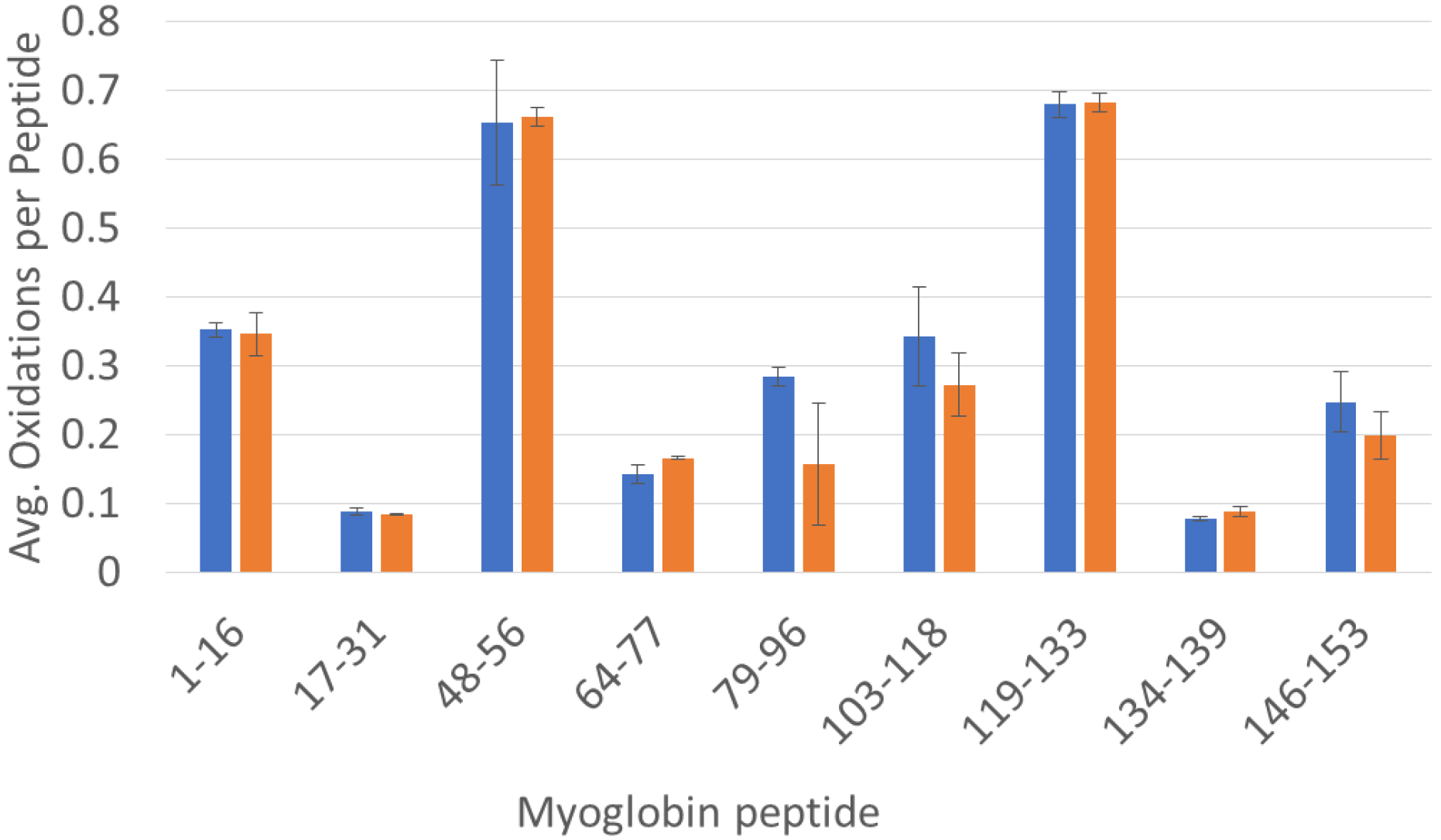
Real time normalization of myoglobin oxidation by inline adenine dosimetry. **(Blue)**Myoglobin oxidized in phosphate buffer at 105 mJ/pulse yielded a ΔAbs_265nm_ of 13.93 ± 2.37 mAU. **(Orange)**Myoglobin oxidized in MES buffer; laser energy was adjusted until an equivalent ∆Abs_265nm_ was achieved (13.27 ± 0.29 mAU) at 130 mJ/pulse. No peptide yielded statistically significant differences in oxidation between samples (α = 0.05).

## Conclusion

HRPF by FPOP is growing in popularity and acceptance, and is recently being used in the biopharmaceutical industry to support protein pharmaceutical development^*40, 41*^. FPOP also has been explored for its uses in a regulatory environment in demonstrating higher order structure equivalence in biopharmaceuticals^*42*^. As FPOP disseminates to a wider audience, the need for convenient and reliable methods for normalizing experiments are necessary in order to allow for reproducibility as well as for extending the technology to the comparison of systems with widely differing radical scavenging properties (e.g. comparison of the effects of excipients on higher order structure in protein pharmaceutical formulations). Inline adenine dosimetry offers a real-time means to standardize experimental conditions, as well as to normalize results obtained in different scavenging environments. We demonstrated that this normalization allows us to obtain identical footprints for proteins in widely differing scavenging backgrounds, which is key for comparability studies. Incorporation of a pinhole aperture as previously reported^*36*^ allows for investigators to change fluence (and, therefore, the amount of radicals generated) without need of correcting for altered beam area by changing the flow rate.

Therefore, with inline adenine dosimetry, experiments could be normalized simply by changing the laser pulse energy at a given repetition rate. Indeed, with appropriate software control, this change can be made automatic based on feedback from the inline adenine dosimeter. When coupled with automated sample collection to ensure that only samples with an equivalent effective hydroxyl radical dose are collected, inline adenine radical dosimetry represents a key innovation towards enabling fully automated HRPF. While our results using Tris buffer demonstrate the need to be careful regarding the UV absorbance of buffer additives, the near linear positive UV response of Tris indicates that it may be possible to use Tris as a positive UV-responsive hydroxyl radical dosimeter in cases where Tris buffer is required.

## Supporting Information

Anomalous adenine dosimetry readings from oxidation of Tris buffer.

## Acknowledgements

S.K.M., R.W.E., J.J.P. and S.R.W. acknowledge research funding from the National Institute of General Medical Sciences (R43GM125420-01) to support commercial development of a benchtop FPOP device. S.K.M. and J.S.S. acknowledge research funding from the National Institute of General Medical Sciences (R01GM127267) for the development of normalization protocols for high-energy FPOP.

